# Tardigrade-Derived Strategy for Low-Cost Storage of Cell-Free Expression Lysates

**DOI:** 10.64898/2026.03.29.715078

**Authors:** Marten Meckelburg, Imre Banlaki, Aukse Gaizauskaite, Henrike Niederholtmeyer

## Abstract

Cell-free expression systems (CFES) are increasingly used alongside conventional biotechnological approaches to accelerate early-stage prototyping and are particularly valuable in point-of-use settings. However, their broader adoption remains limited by time- and cost-intensive preparation, as well as stringent cryogenic storage requirements. To address this, several studies have explored lyophilization with protective additives to generate stable, solid-state CFES. These approaches had to balance the protection gained with a loss of activity due to the additives. In this study, we present a CFES that contains a tardigrade-derived Cytosolic-Abundant Heat-Soluble (CAHS) protein to protect the biosynthetic machinery in lysates from damages during drying. We show that the CAHS protein, without any other additives, preserves protein synthesis activity during low-cost room temperature desiccation, while unprotected lysates are affected in mRNA synthesis kinetics and translation yields. The diversity of tardigrade-derived protective proteins is a treasure trove for cell-free synthetic biology, in particular for making CFES more accessible and portable.

**Graphical abstract:** 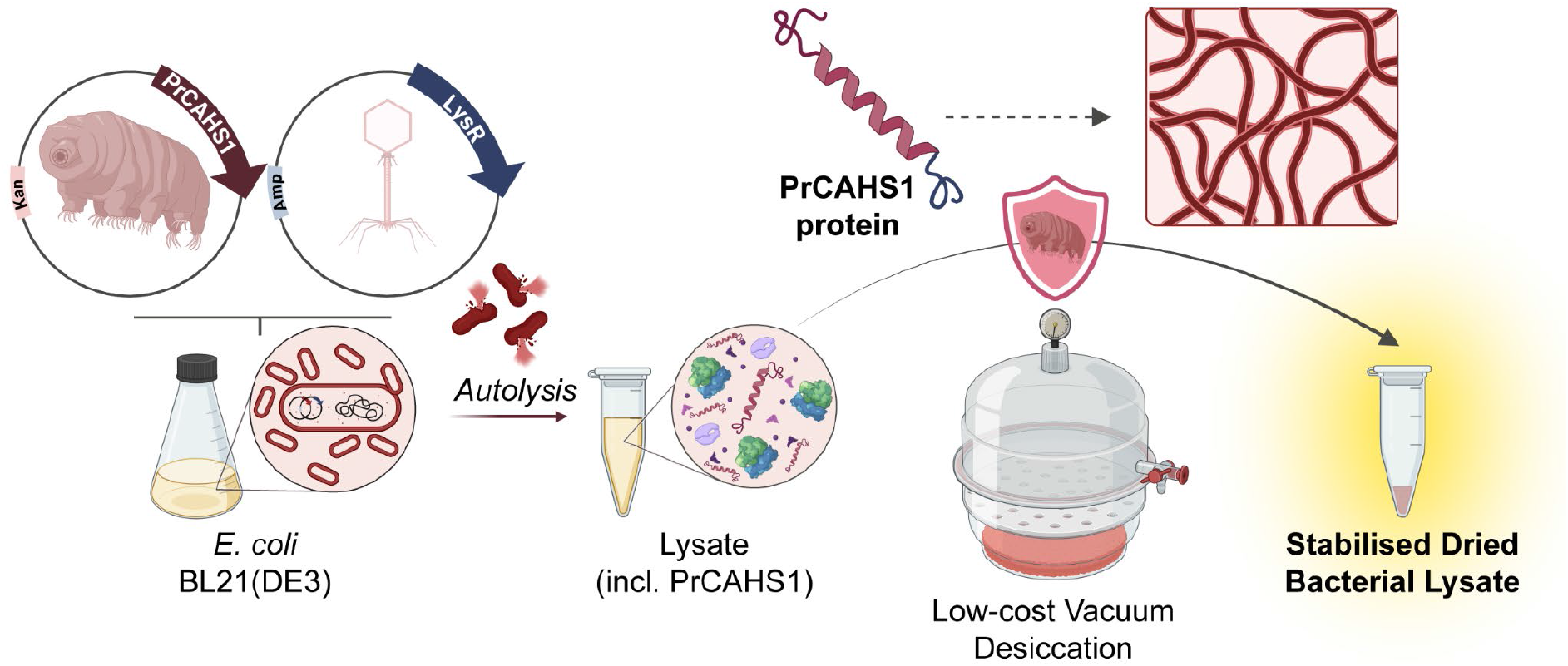

## Introduction

Climate change has incentivised the search for sustainable bio-based approaches for the synthesis of high-value compounds and pharmaceuticals, instead of the traditional chemical production from fossil hydrocarbons. While traditional approaches employ living cell-factories, cell-free expression systems (CFES) have gained prominence owing to their improved reaction control and ability to circumvent major limitations such as cell-viability and lengthy engineering cycles^1,2^. Versatility, speed, and the open reaction format have made CFES indispensable for synthetic biology. *Escherichia coli* lysate-based CFES are widely used to rapidly synthesize protein or peptide variants, to prototype synthetic gene circuits and in point of care applications such as biosensing^3–5^. They are easy to use with standard laboratory methods as well as in automated setups. Cell-free reactions are particularly attractive in field applications and educational settings because they bypass the biosafety and regulatory concerns associated with engineered cells, give rapid outputs and usually do not require expensive equipment^6–10^. In addition, portable CFES have even been proposed for on-demand production of biopharmaceuticals^11,12^.

Despite high interest, the widespread adoption of CFES is limited by their requirement of −80 °C storage and cold-chain logistics, restricting access for underfunded labs and field applications in resource-limited settings^6,13^. State-of-the-art strategies to stabilize CFES for low-cost storage and distribution rely mainly on lyophilization. Lyophilization is often combined with protective additives, particularly sugars (trehalose, sucrose, lactose, maltodextrin, dextran), polyols, and multi-component formulations tailored to a specific extract type^13–16^. These approaches have been shown to alter CFES metabolism and therefore require extensive, system-specific optimization, demanding special equipment and costly resources^13,15,17^. This underscores the need for low-cost, easily integrated, protective strategies. In previous work, Guzman-Chavez et. al. (2022) explored the alternative production of low-cost CFES by simply drying lysate in a conventional vacuum desiccator instead of lyophilisation^13^. Even though their results showed that damage occurring during vacuum desiccation tends to be more severe than lyophilisation, the retained activity encouraged us to investigate further ways to improve desiccation and expand the production repertoire of low-cost, portable CFES^13,15^.

In the spirit of synthetic biology, we looked to solutions found in nature for surviving desiccation. Probably the most prominent organisms known to survive a host of adverse conditions are tardigrades^18,19^. There has been much interest from fundamental and applied biology alike to find out how tardigrades exhibit these exceptional capacities and whether they can be transferred to other biological systems.

Such transferability was successfully showcased for protection of biologicals against environmental challenges like osmotic stress^20,21^, radiation^22,23^ and chemical stressors such as mutagens^24^ or reactive oxygen species^25^. Synthetic biologists, for instance, heterologously expressed tardigrade proteins in human cells to protect them from chemically induced apoptosis^26^. In another work, the transfection with mRNA encoding a tardigrade protein protected DNA in mammalian tissue against radiation damage^23^.

We were inspired by tardigrades’ remarkable desiccation tolerance. To survive extreme water loss, tardigrades enter a vitrified “tun” state, where the organism’s metabolism comes to a complete stand-still^18,19^. Multiple mechanisms protect tardigrades from desiccation-related damage. Among them is the production of several classes of intrinsically disordered proteins such as secretory-, mitochondrial- and cytoplasmic-abundant heat soluble (SAHS, MAHS, CAHS) proteins^19,27^. Since their discovery, these proteins have found applications in the protection of various biologicals. For example, the external application of purified SAHS proteins protected liposomes and bacteria during desiccation^28^. In other studies, CAHS proteins have been used for the protection of purified enzymes such as lactate dehydrogenase^27,29–31^, but also for the reliable mitigation of desiccation related damages in whole cells and giant unilamellar vesicles^21,27,32–34^. CAHS proteins form protective gel-like fibrous networks upon desiccation (Figure S1A)^35,36^. Once formed, these networks are believed to mitigate desiccation related damages through a combination of different mechanisms such as the formation of hydrogen bond networks that substitute the loss of the hydration shell, and restricting protein mobility as well as conformational changes^30,37^.

Particularly relevant to this work is a study by Boothby *et al*.^38^ which demonstrated that multiple CAHS proteins significantly increased microbial survival rates during desiccation. We selected two proteins, CAHS107838 (UniProt ID P0CU51, referred to as PrCAHS1) and the close homolog CAHS106094 (UniProt ID P0CU52, referred to as PrCAHS2) from *Paramacrobiotus richtersi*, that performed best in improving *E. coli* survival, in order to test their ability to protect bacterial lysates from desiccation damages, and for creating stable, stolid-state CFES. We show that PrCAHS1, produced in the *E. coli* strain for lysate production, preserves protein-synthesis capacity of low-cost, room-temperature desiccated lysates. We monitored mRNA and protein synthesis dynamics in PrCAHS1-protected and control lysates and show that drying of unprotected lysates mainly damages the translation machinery. Based on protein structure predictions and experiments with the desolvating agent 2,2,2-trifluoroethanol (TFE), we propose that PrCAHS1 forms protective higher order protein assemblies during desiccation, similar to other CAHS proteins that are known to form gels and filamentous networks^31,35,39^.

## Results and Discussion

### Integration of CAHS protein is compatible with existing lysate preparation workflows

For lysate preparation, we made use of the autolysis method developed by Didovyk *et al*.^40^, where an autolysis plasmid produces λ-phage endolysin to enable cell lysis by simply freeze-thawing the biomass. To produce lysates enriched with CAHS protein, the lysate production strain *E. coli* BL21(DE3) was transformed with a second plasmid carrying the CAHS gene under control of the inducible T7 promoter (Figure 1A, Figure S2).

**Figure 1.**
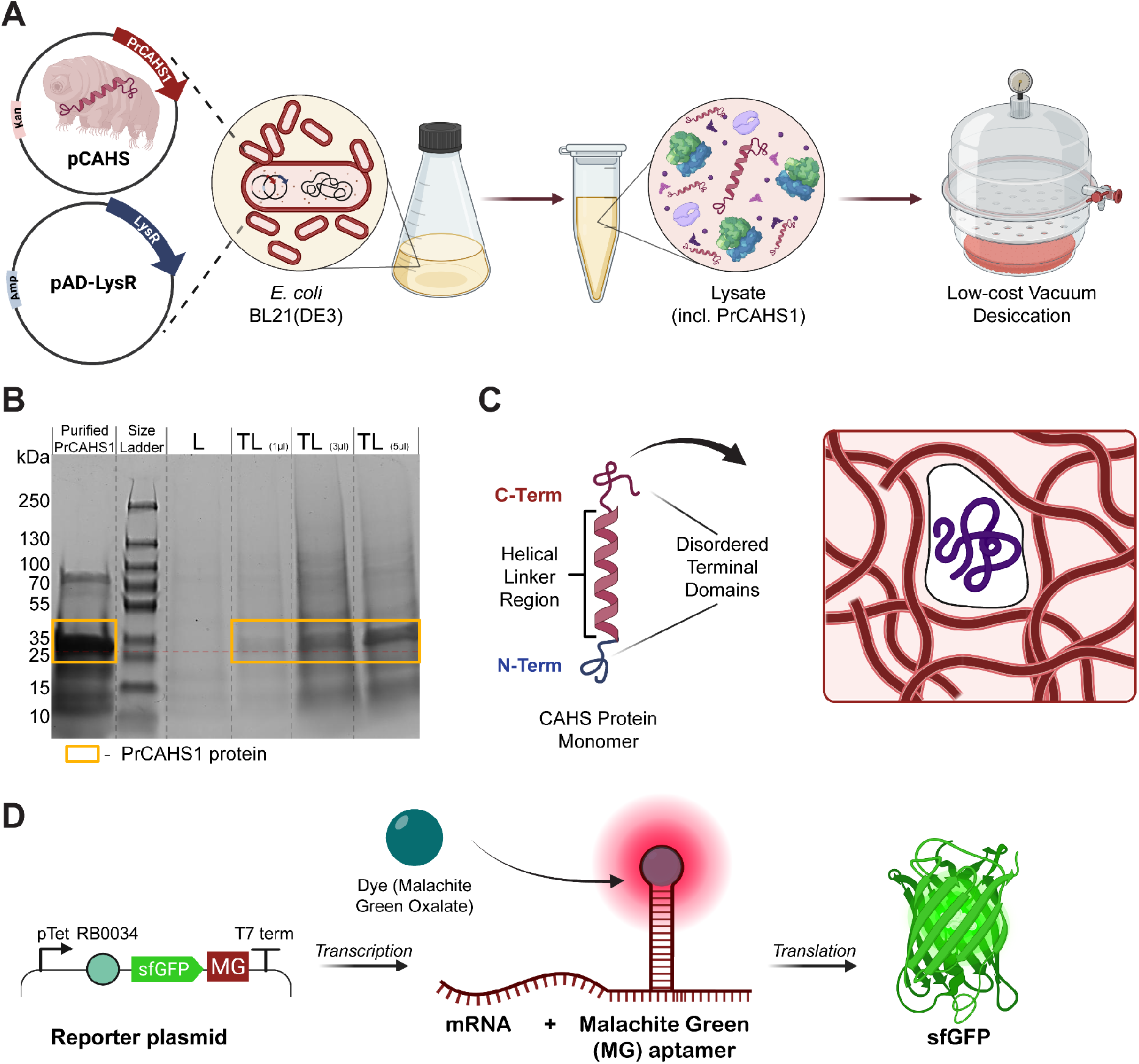
Conceptual design of the study and generation of PrCAHS1 protein containing lysates. **(A)** Schematic of the workflow to produce solid-state lysates. A plasmid encoding for the inducible expression of PrCAHS1 protein is co-transformed with the constitutively expressing autolysis plasmid^40^ into *E*.*coli* BL21(DE3). During cultivation, the expression of the CAHS protein is induced. The biomass is processed into lysate containing CAHS protein variants and is subjected to desiccation in a conventional vacuum dessicator. **(B)** 12 % SDS page showing the presence of PrCAHS1 protein in the final lysate. Lanes (left to right) contain: purified PrCAHS1 (5 µl, 50.82 µg); protein standard; “L” - lysate produced from BL21(DE3) strains lacking CAHS protein (3 µl); “TL” - lysate with PrCAHS1 protein (added in 1, 3, 5 µl quantities). Note: all lysate samples were boiled at 95 °C and centrifuged prior to processing for the gel. Expected size of PrCAHS1 = 26.5 kDa (yellow box) **(C)** Schematic of the structure of CAHS monomers and their proposed transition into a mesh-like network that stabilizes client proteins enclosed within. **(D)** Schematic illustration of the function of the transcription and translation reporter plasmid. Transcription is monitored in the red spectrum by the MGO-aptamer complex. Translation is monitored by the green signal of synthesized sfGFP.

Before evaluating CAHS-mediated desiccation protection, we first confirmed that CAHS proteins were soluble and detectable in the *E. coli* lysates. To avoid altering protein function, we did not add an affinity tag. However, the presence of CAHS proteins is easily verified due to their heat solubility^41^. To check for the presence of CAHS proteins in the lysates, we boiled and centrifuged aliquots, before using the supernatant for an SDS-PAGE. In the standard lysate (L), without CAHS protein, little protein remained after the boiling. Conversely, we detected a clear band corresponding to PrCAHS1 in the tardigrade lysate (TL) (Figure 1B). In addition to a band corresponding to PrCAHS1, boiled TL samples showed additional protein bands on the gel. This observation may suggest that PrCAHS1 helps other proteins, in the lysate, remain in solution during boiling and hints at interactions of heat soluble PrCAHS1 and cytosolic protein at high temperatures. Small heat shock proteins are used by tardigrades to achieve extreme heat resistance, but also show potential roles in desiccation tolerance so this effect may hint to a conserved dual role in CAHS proteins as well^42^. This suspicion is supported by another study where two CAHS variants showed some protective effects on lactate dehydrogenase under heat-stress^29^.

Due to its insolubility upon heterologous expression, PrCAHS2 was disqualified for the production of cell-free expression lysates (Figure S3). While PrCAHS2 can be resolubilized by boiling, it is not compatible with our lysate preparation workflow.

The rationale behind our design, combining autolysis with protective CAHS-protein production, was that such a lysate can be produced without any specialized equipment or processing other than a liquid culture, a freezer, a vortex and a desiccator. We hypothesized that during desiccation of a PrCAHS1-containing tardigrade lysate (TL), the concentration of the CAHS protein will increase to the point where a matrix will form around sensitive components and protect the lysate (Figure 1C). This stands in contrast to traditional means of production, where lysates are prepared by sonication or mechanical lysis^43,44^, and protective sugars and polymers have to be added subsequently before lyophilization^13–16^.

To assess the stability of lysates against desiccation-related damages, we designed a dual reporter plasmid to be expressed in our CFES (Figure 1D). With this reporter, translation was quantified by super-folder green fluorescent protein (sfGFP) synthesis, whereas transcription was measured via the fluorescent malachite green aptamer (MG), which we integrated in the 3’ end of the reporter mRNA transcript. By binding malachite green oxalate (MGO), added to the CFES reaction, the MGO-aptamer complex becomes fluorescent in the red spectrum (emission measured at 656 nm). As established in other studies, this allowed us to simultaneously monitor mRNA and protein abundance^45,46^. Expression of the reporter is controlled by a strong promoter (pTet) paired with a strong ribosome binding sequence (RB0034) (Figure 1D).

### Intrinsic expression of *PrCAHS1* reduces desiccation damages

To test the protective capacity of PrCAHS1, we produced lysates with (TL) and without the protein (L) and used two storage conditions. As a bench mark, liquid aliquots of each lysate were stored in the −70 °C freezer (standard conditions, or “fresh”). Solid-state aliquots, from the same lysate batches, were produced by room temperature desiccation under vacuum and in the presence of a silica desiccant (Figure 1A). These dry samples were packed in a nitrogen atmosphere and vacuum sealed including a silica desiccant sachet (Figure S4). To store the dried aliquots, we placed them in a closed styrofoam box and kept them at room temperature for a week or up to a month.

After a week of storage, dry aliquots were rehydrated by adding water to the original volume of the sample. We compared fresh lysate samples with rehydrated dry aliquots by measuring their transcription and translation activity in the plate reader (Figure 2A). The same energy buffer and reporter plasmid were added to all lysate samples. We observed the typical CFES protein production kinetics, with sfGFP increasing initially and plateauing, starting at 3 h, for both control and tardigrade lysates, as well as for fresh and desiccated lysates (Figure 2A). While sfGFP production curves were markedly lower for dried lysates without the PrCAHS1 protein (L), PrCAHS1-containing lysate activity was not notably affected by drying. As the PrCAHS1-containing TL lysate had a lower activity in its fresh state, we normalized dried to fresh sfGFP yields (Figure 2B, C). Each plate reader run was conducted as three technical replicates per lysate type and treatment.

**Figure 2.**
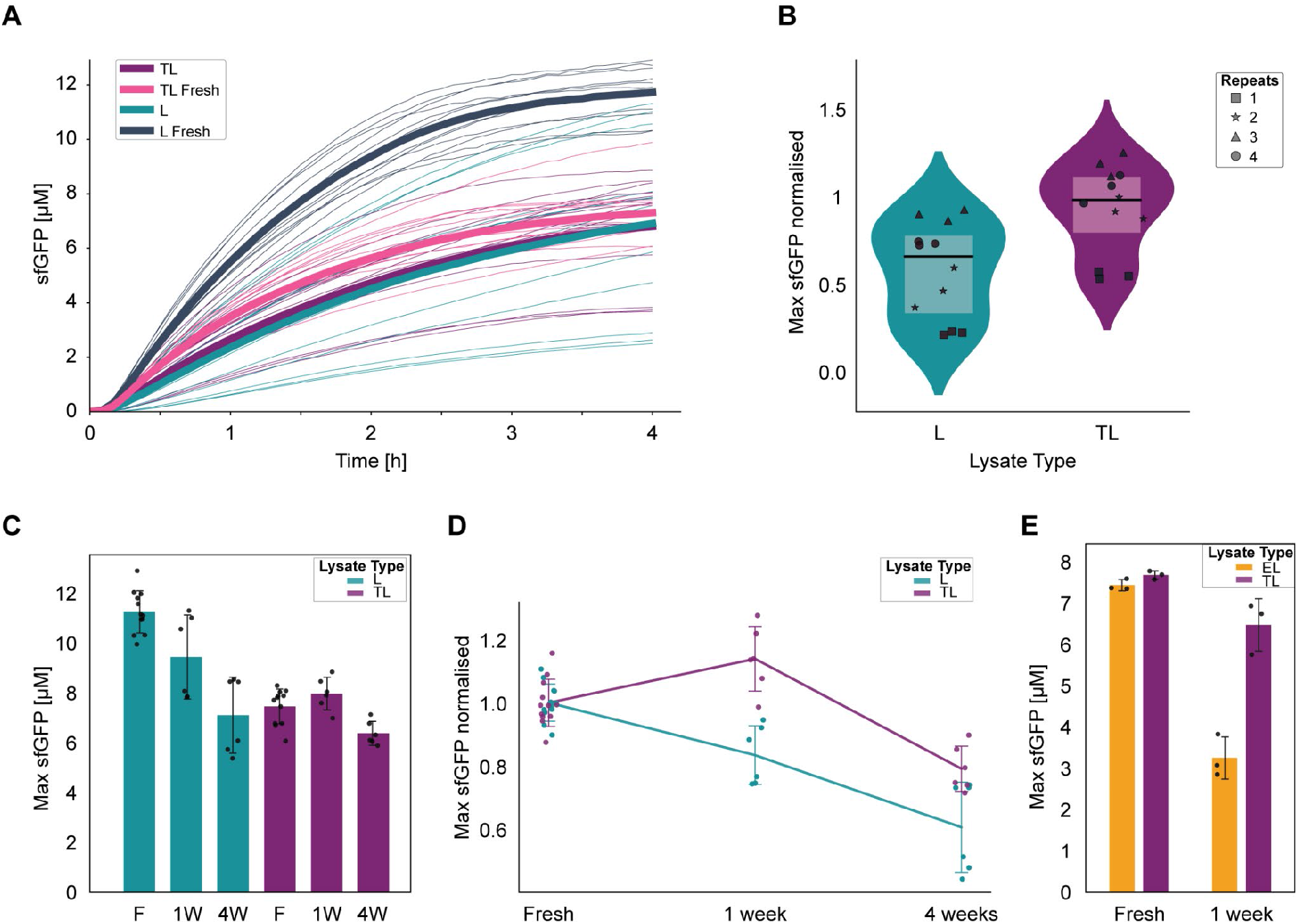
PrCAHS1 protein preserves translation yields from immediate, desiccation-related damages. **(A)** sfGFP synthesis kinetics for different lysate types (L and TL) and treatments (desiccated and fresh). Solid thick lines show mean values and thinner lines show individual replicates from four short-term desiccation experiments (n = 12, across four experiments with n = 3 technical replicates respectively). **(B)** Violin plots of normalized maximum sfGFP concentrations in dried lysates (L and TL) after 4 h. Symbols distinguish individual experiments (n = 12, across four experiments with n = 3 technical replicates respectively). The line within the violin plot represents the median, upper and lower border of the box plots representing upper and lower quartiles respectively. **(C)** Final sfGFP yields (maximum at 4 h) at fresh, 1-week, and 4-week time points. Bars show means ± SD (n = 6; fresh n = 12). Individual data points are overlaid. **(D)** Normalized sfGFP yields as a function of storage time. Lines show means ± SD (n = 6; fresh n = 12). **(E)** Final sfGFP yields of EL and TL, fresh and 1 week after desiccation. Bars show means ± SD (n = 3 technical replicates). Abbreviations: Standard Lysate (L), produced from a strain containing only the autolysis plasmid; Tardigrade Lysate (TL), produced from a strain expressing PrCAHS1 protein; Empty vector Lysate (EL), produced from a strain carrying the autolysis plasmid plus the empty expression vector.

Comparing different, independent desiccation experiments, we found that inflicted damages can vary widely with retained activity ranging from 10-75 % compared to fresh lysate (Figure 2B). Yet, across all desiccation runs, TL consistently retained higher activity than L (Figure 2B, Figure S5). In some of the experiments, the rehydrated TL showed even higher translational activity than the fresh equivalent resulting in a normalized fluorescence larger than 1 (Figure 2B). This could potentially be due to the vigorous mixing required to redissolve the desiccated material. If the activity was close to unaffected, the additional dissolved oxygen could boost the activity beyond the fresh lysate, as dissolved oxygen was shown in previous publications to be a limiting factor of aerobic CFES^47,48^.

When looking at activity trends over longer storage times we observed an initial strong protective effect in one week old TL samples but a decrease in activity, following the trend in L, when testing four week old aliquots (Figure 2D). Due to the initial protection however, the four week old TL samples still had higher normalized activity than the unprotected L samples. These results lead us to conclude that the PrCAHS1 protein protects the lysate components during desiccation but does not significantly protect them from chronic long-term degradation during storage. This is reminiscent of results from another study where CAHS proteins (CAHS3 and MAHS) protected yeast from acute osmotic stress but not from chronic osmotic stress^21^. To enable extended storage, future work could explore storage of desiccated lysates at lower temperatures such as 4 °C or −20 °C, which are more widely available and economical than cryogenic storage. As PrCAHS1-protected lysates are stable at room temperature for a week, reagents can be shipped without cooling, while long-term storage might benefit from decreased temperatures.

Comparing the tardigrade lysate TL to the unaltered lysate L, we found that fresh TL only reached about 60 % of activity (Figure 2A, C). The reduced activity of fresh TL lysates might be due to the IPTG induction of T7 RNA polymerase and the synthesis of PrCAHS1 protein, as well as the addition of a second antibiotic during biomass growth. To rule out that activity conservation in TL was an artefact of the decreased overall activity of TL samples, which potentially masked damages accrued during desiccation, we produced a lysate (EL) from an empty vector control strain. EL differed from L by addition of an empty plasmid containing the second antibiotic resistance gene. Indeed, fresh EL had a similar activity to TL suggesting the decreased activity was due to the additional plasmid and IPTG induction. With these lysate pairs, the protective effect of PrCAHS1 could be confirmed as the EL lost more than 50 % activity after one week of storage while the TL retained more than 80 % of its activity (Figure 2E). As EL and TL had comparable activities in the fresh state, we conclude that the presence of the protective PrCAHS1 protein had no measurable effect on lysate activity. Instead, it is more likely that the presence of a second antibiotic and IPTG-induction of the T7 RNAP reduced lysate activity by about 40 %^49^. In future work, this could be addressed by combining autolysis and PrCAHS1-production on a single plasmid, and exchanging the T7 promoter for an *E. coli* RNA polymerase promoter.

### Desiccation damages affect dynamics of transcription and translation

In addition to testing overall protection by CAHS protein, we were also interested in specific effects on transcription and translation (Figure 3). As mentioned, the reporter plasmid coded for sfGFP and a Malachite Green aptamer (MG) on a single mRNA. This allowed us to monitor transcriptional activity, from the MG signal, independently of translation via sfGFP synthesis. Comparing all conditions, we observed two qualitative modes of aptamer signal kinetics (Figure 3A). After an initial increase of signal, fresh and undamaged, rehydrated TL samples started to decrease in signal around the 2h mark. In contrast, damaged lysates did not show a peak in their fluorescence, instead, they increased steadily in signal or plateaued. This trend is highlighted when we compare the difference in aptamer signal (ΔMG) between the 2 h and 4h time points (Figure 3B). While fresh and protected TL samples show values in the negative range, unprotected lysate samples exhibit mostly positive values.

**Figure 3.**
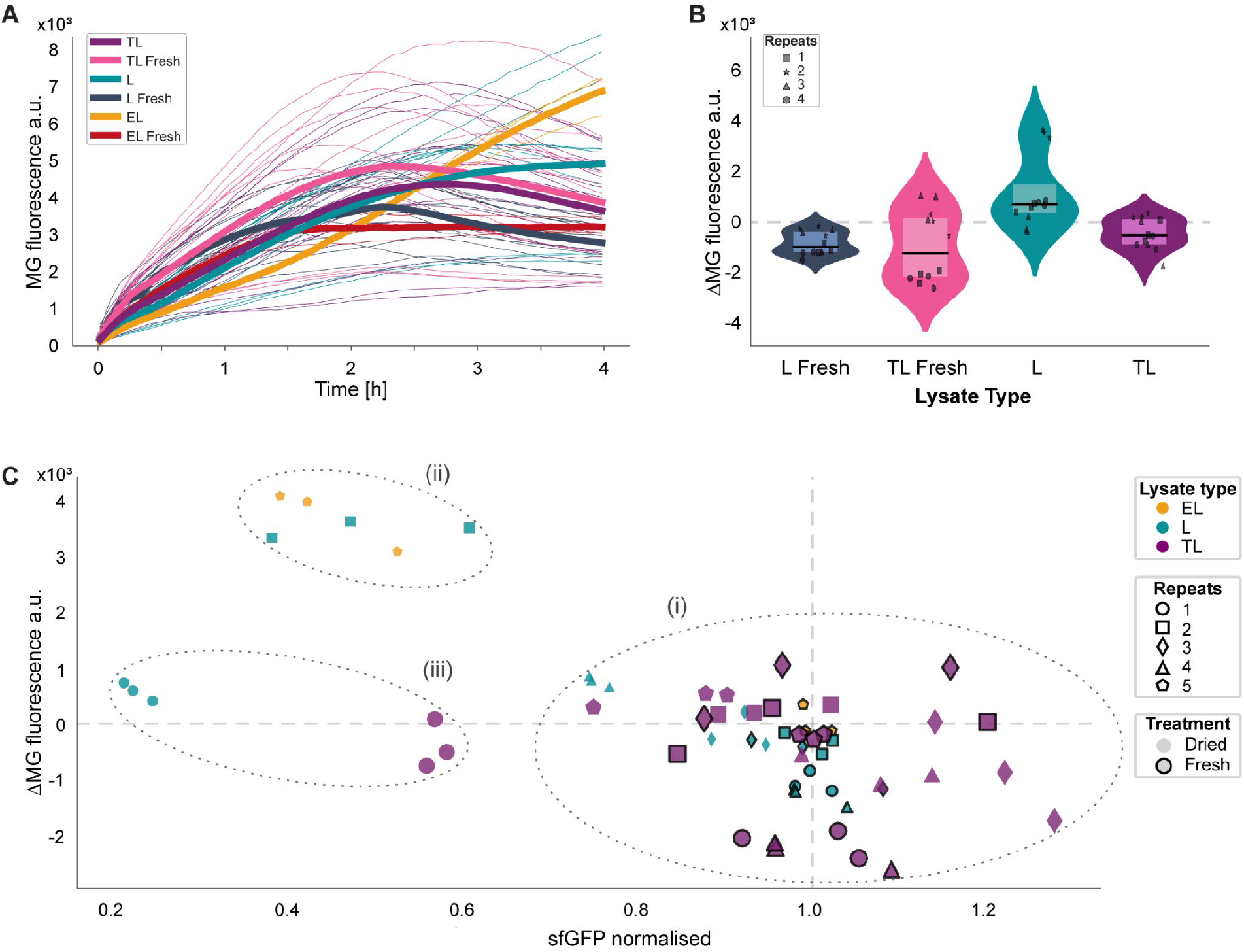
Effects of PrCAHS1 protein on transcriptional dynamics after desiccation. **(A)** Malachite green (MG) fluorescence kinetics across four separate short-term (1 week) desiccation experiments of different lysate types (EL, L, TL) compared to their fresh controls. Solid thick lines show mean values and thin lines correspond to individual replicates from four short-term desiccation experiments (n = 12 per group, except EL (n = 3)). **(B)** Violin plot comparing the change of malachite green fluorescence (ΔMG) between 2h and 4h across four separate short term desiccation experiments (with n=3 technical replicates each). Symbols distinguish individual desiccation runs. The line within the violin plot represents the median, upper and lower border of the box plots representing upper and lower quartiles respectively. ΔMG=0 is indicated with a dashed line. **(C)** Scatter plot visualizing tradeoffs between sfGFP production and the change of malachite green fluorescence (ΔMG) between 2h and 4h. (i) Cluster of fresh and lightly damaged samples; (ii) cluster of samples with medium damage; (iii) cluster of heavily damaged samples. Symbols distinguish individual desiccation experiments; colors indicate lysate identity; fresh lysates are indicated with black frames around datapoints. Abbreviations: Standard Lysate (L), produced from a strain containing only the autolysis plasmid; Tardigrade Lysate (TL), produced from a strain expressing a PrCAHS1 protein; Empty vector Lysate (EL), produced from a strain carrying the empty expression vector plus autolysis plasmid.

Prior work suggests that in high performing CFES, transcription and translation are in resource competition, with translation outcompeting transcription when sufficient mRNA is present^46^. With the transition to a translation dominated regime, and parallel mRNA degradation, we can explain the MG signal decrease in highly active CFES. The loss of this competition in unprotected, dried lysates suggests that desiccation more strongly affects the translational machinery. This notion is even more apparent when we examine the different desiccation experiments individually and compare ΔMG to normalized sfGFP production (Figure 3C, Figure S6). In experiments where little damage was inflicted overall, dried samples exhibited similar negative ΔMG as fresh controls (Figure 3C (i)). At intermediate damages, ΔMG becomes positive in unprotected lysates but PrCAHS1 protects the lysate from significant loss of translational activity (Figure 3C (ii)). However, under heavy damage conditions (desiccation repeat no. 1), even transcription is affected. When a lysate is heavily damaged, ΔMG is close to zero due to low overall MG signal, and sfGFP production is severely reduced (Figure 3C (iii), Figure S6).

Comparing MG signals of fresh and rehydrated samples during the initial phase of the reaction, we did not notice significant differences between PrCAHS1-protected lysates and unprotected lysates. In general, MG signals of all desiccated samples, regardless of PrCAHS1, increased at slightly lower rates than in their fresh counterparts (Figure 3A, Figure S7). As the exact interactions of PrCAHS1 with cytosolic components are unknown, we can only speculate on the reason for the difference in protection between transcription and translation machinery. However, evaluating the different experiments with unprotected lysate, it appears that RNA polymerase is less affected by desiccation overall, compared to the translation machinery. While protein yields of a PrCAHS1-protected CFES are hardly affected by the slightly reduced initial mRNA synthesis rates, unprotected lysate with comparable initial mRNA synthesis is severely impacted by desiccation in its potential for translation.

### TFE supplementation induces the formation of higher order assemblies of CAHS proteins

To better understand the potential action of CAHS protein in our lysates, we purified PrCAHS1 and PrCAHS2, by taking advantage of their heat solubility properties (see Methods and Figure S8)^41^. In previous studies it was shown that many CAHS proteins consist of a polyampholytic helical linker region and largely disordered terminal domains (Figure S1A). At high concentrations and under desiccation, these proteins form higher order structures such as fibrils and gels^35,36^. This could be in a process analogous to the latest model for *Hypsibius exemplaris* CAHS-D assembly, where antiparallel dimers form through the polyampholytic, amphipathic character of the helical linker, which mediates contacts between the α-helical faces of the two chains^36^. These dimers then polymerise to form fibrils which can further assemble into higher order fibers. Notably, our AlphaFold2 prediction for the PrCAHS1 protein structure showed formation of a similar antiparallel dimer (Figure S9). An alternative model proposes the assembly of CAHS-D dimers in a concentration-dependent manner into supercoiled 22-mer bundles, which polymerize through end-to-end interactions to form a fibrous gel (Figure S1)^35^. In contrast to CAHS-D, concentrated protein solutions of up to 13 mg/mL were not gel-like or notably viscous for PrCAHS1 and PrCAHS2. Indeed, previous work showed that different CAHS variants vary in their propensities to form higher order structures^32^.

In order to mimic desiccation conditions, previous studies have supplemented the desolvating agent 2,2,2-trifluoroethanol (TFE) to CAHS protein solutions^32,50,51^. TFE displaces water from protein surfaces, which promotes intramolecular hydrogen bonds and electrostatic interactions within the polypeptide, resulting in an ordered secondary structure that may approximate the “dry” protein structure under desiccation conditions (Figure 4A)^52^. TFE-induced structural changes heavily depend on both protein and TFE concentrations^51^. Therefore, the TFE assays used in this study serve as a purely qualitative assessment of response to the desolvating agent and may not precisely reflect processes occurring during the desiccation process.

**Figure 4.**
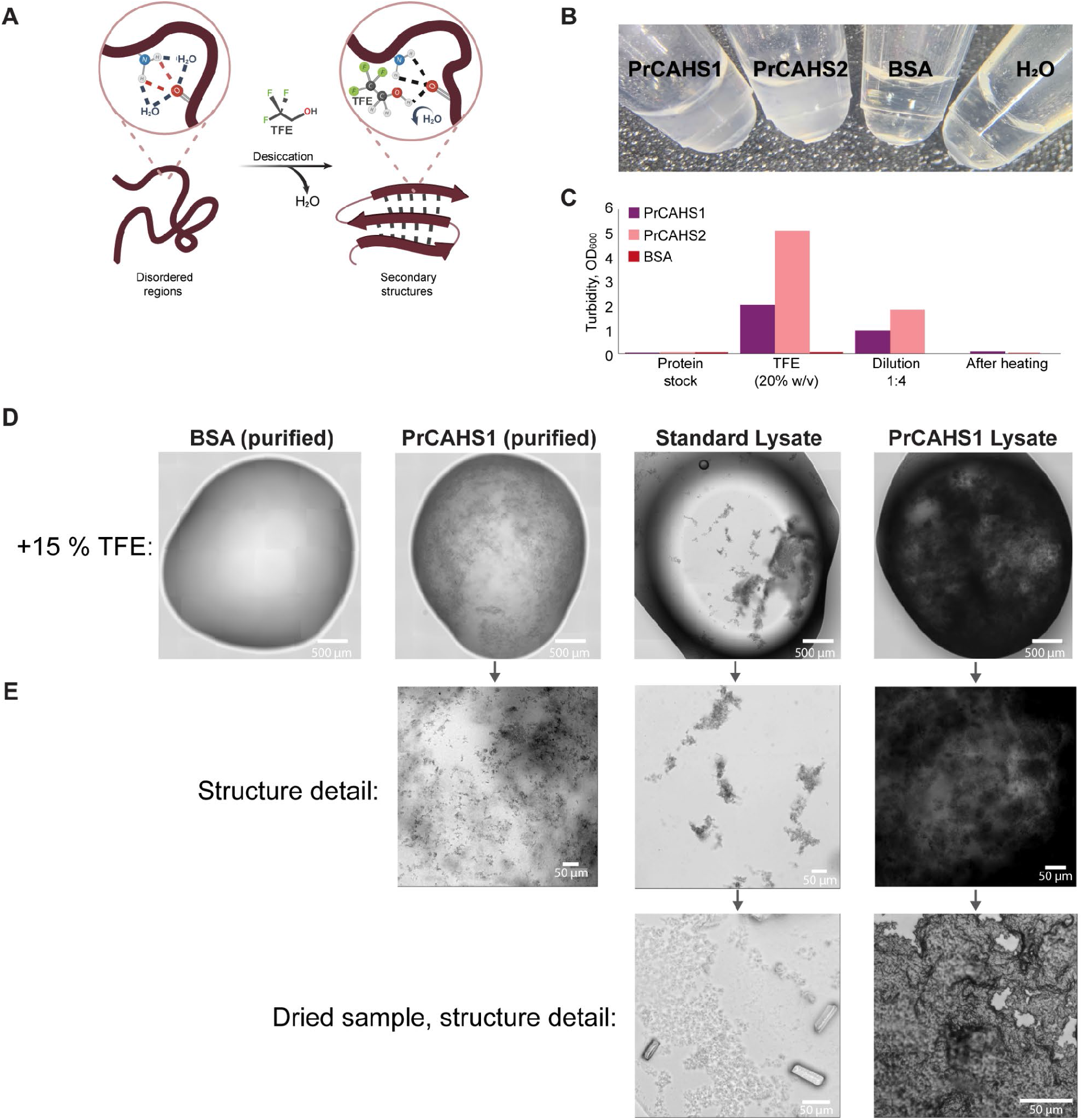
Formation of higher order assemblies of CAHS proteins, induced by trifluoroethanol (TFE) **(A)** Schematic illustrating how TFE promotes secondary structure formation and higher-order assembly in CAHS proteins^32,51,52^. In aqueous solution, water weakens intramolecular hydrogen bonds by competition. TFE addition mimics the effects of water removal during desiccation through water displacement, promoting intramolecular hydrogen bonding and secondary structure formation in otherwise disordered regions. **(B)** PCR tubes containing 10 mg/mL protein solutions after TFE supplementation (20 % v/v). PrCAHS1 and PrCAHS2 = purified tardigrade proteins + TFE; BSA = Bovine Serum Albumin + TFE; H_2_O = water + TFE. **(C)** Turbidity (OD_600_) measurements of the samples in (B) under different conditions: prior to TFE addition (protein stock), after TFE addition (20 % v/v), after 1:4 dilution with water, and following heat treatment at 95 °C post-TFE addition. **(D)** Brightfield microscopy images of TFE-induced (15 % v/v) assemblies. From the left: purified BSA, purified PrCAHS1 protein (10 mg/mL), standard lysate (boiled and centrifuged) and lysate containing pre-expressed PrCAHS1 proteins (boiled and centrifuged). Scale bars 500 µm. **(E)** Magnified structure details of samples from section (D). The bottom row shows structure details of dried lysates after water and TFE evaporation. Scale bars 50 µm.

To characterize PrCAHS1 and PrCAHS2 further, we assessed the formation of higher order assemblies in purified protein samples and in lysate samples with or without CAHS proteins, mediated through the supplementation of TFE (Figure 4).

After adding 20 % TFE, solutions of 10 mg/mL PrCAHS1 and PrCAHS2 turned highly turbid, suggesting the assembly of structures that scatter light, while a bovine serum albumin control sample remained clear (Figure 4B). To rule out the formation of irreversible aggregates and covalent crosslinks, we heated TFE-treated PrCAHS1 and PrCAHS2 to 95 °C prior to a second optical density measurement (Figure 4C). As expected^35^, heating the CAHS proteins reduced turbidity to values comparable to BSA controls and samples before treatment with TFE (Figure 4C).

In a repeat experiment, we examined 15 % TFE-treated samples under the microscope. Purified PrCAHS1 appeared to form dense, fibrous structures, whereas the BSA control remained unaffected (Figure 4D). We also compared lysates containing intrinsically expressed PrCAHS1 with the control lysate without the tardigrade protein. As lysates contain high concentrations of all *E. coli* cytoplasmic proteins, which obscured our results by aggregating, we boiled the two lysate types, and removed denatured protein by centrifugation. As we know from gel analyses (Figure 1B), boiled PrCAHS1-lysates contain some additional leftover proteins. These clarified, boiled lysates were then treated with TFE to assess whether PrCAHS1, directly derived from the CFES, is also able to form higher order assemblies. While the control lysate sample, without PrCAHS1, remained mostly clear except for a few larger aggregates, the PrCAHS1-containing lysate turned opaque with fibrous structures (Figure 4D, 4E). After leaving these samples to air dry overnight, the dried PrCAHS1-containing sample formed a continuous solid, whereas the standard lysate left only little residue (Figure 4E, bottom row). This observation fits models for CAHS protein-mediated desiccation resistance. A tight network-matrix is believed to protect client proteins by restricting their conformational freedom, substituting the loss of hydrogen bonding, or coordinating residual water while shielding clients from reactive species and aggregating partners^35,37^.

Although the exact assembly mechanism of PrCAHS1 and its interactions with cytosolic proteins during desiccation are beyond the scope of this work, we observed clear qualitative effects when exposing PrCAHS1 samples to the desolvating agent TFE and to heat. The behavior of PrCAHS samples, in response to TFE, points to the formation of higher order assemblies (Figure 4). We also have indications for interactions with cytosolic proteins because some additional proteins are protected from aggregating during boiling (Figure 1B).

The mechanism of desiccation protection has been shown to vary between CAHS variants. For example, a recent study showed that CAHS variants with a reduced tendency to form higher-order assemblies were more effective at protecting certain enzymes, such as citrate isomerase. Other variants, which assemble more readily, provided superior protection for others, including lactate dehydrogenase^31^. Furthermore, studies comparing different CAHS proteins identified homologs forming filamentous networks or spherical structures^32^. These findings suggest a mechanistic diversity in CAHS protein assembly and protection, opening up the prospect of combining different CAHS proteins in order to synergize their protective effects on CFES, potentially against chronic desiccation damages as well.

## Conclusion

Cell-free protein expression has become an indispensable technology in synthetic biology, biotechnology and fundamental research alike. With prohibitive costs and specialized know-how limiting the use of CFES, much work has been dedicated to facilitate and simplify its deployment. With our study, we supported this effort by integrating desiccation protection in biomass production, removing a step in the standard protocol for solid-state lysate production.

We have shown that PrCAHS1 containing lysates are a viable, protein-based solution to protect CFES from desiccation damages. Lysates protected by tardigrade CAHS protein do not require chemical preservatives, thus mitigating metabolic perturbations common in other formulations. Furthermore, the combination with the autolysis system makes production and scale up convenient and easy with standard laboratory equipment.

What we did not address in this work is the storage of energy components. In some applications, energy components are premixed with lysate to create a combined solid state CFES^14^. As our desiccation method is slower than lyophilization and occurs at room temperature, a combined desiccation would suffer from metabolic idling consuming much of the extra energy provided^49,53^. However, as most energy buffer components are procured in dried form, it may be as easy as combining them in a powder mill and adding them after desiccation.

Another limitation we noticed is the chronic degradation occurring during prolonged storage, which proceeded at similar rates in PrCAHS1-protected and unprotected lysates. Technical improvements, such as storage under noble gas, with additional oxygen scavengers, and at lower temperatures, such as 4 °C or −20 °C, may mitigate chronic degradation. Alternatively, synergistic combinations of different CAHS proteins, as is the case in tardigrades, may enable long-term storage in ambient conditions. Indeed, the vast diversity of different proteins, many occurring in the same organisms, may prove a treasure trove for synthetic biologists searching for ways to increase biological tolerance of harsh conditions. Such further investigation may then focus on combining functions, proven in isolation, to elucidate possible cooperative effects.

## Methods

### Plasmid assembly

Plasmids used for Golden Gate assembly were purified using NEB Monarch® Plasmid Miniprep kit and dual reporter plasmid was purified using Macherey-Nagel NucleoBond® Xtra Midiprep kit, both according to the manufacturer’s protocols.

Constructs were generated through Golden Gate assembly utilizing the “Marburg collection”^54^. The codon optimised tardigrade genes were ordered from GenScript (GenScript Biotech Corp.). Both level 0 and level 1 assemblies were conducted in adherence to the instructions provided with the enzyme manufacturer (NEB). Constructs were transformed into DH5α *E*.*coli* strains, verified using colony PCR and sequenced.

### Lysate production

Lysates were generated from *E. coli* BL21(DE3) harbouring an autolysis plasmid, in accordance with the workflow described by Didovyk *et al*.^40^, with some adjustments to improve accessibility and to co-express tardigrade genes during the process. Briefly, competent BL21(DE3) were cotransformed with the autolysis plasmid pAD-LysR (Addgene #99244) and the plasmid for inducible PrCAHS variants expression (TL lysate variant), or the empty vector (EL variant) (Table S1), and grown on an LB agar double selection plates (Ampicillin and Kanamycin). The standard control lysate strain (L) was only transformed with pAD-LysR. To produce biomass for lysis, we seeded overnight cultures in 5 mL 2xYTPG medium (supplemented with glucose at a final concentration of 100 mM) growing them at 37 °C and 200 RPM. These cultures were used to seed larger 800 mL cultures, in 2xYTPG medium, in baffled flasks the following day, using 1 mL as inoculum (37 °C, 200 RPM). Growth was monitored via the measurement of OD_600_ using a spectrophotometer (Implen GmbH). *PrCAHS* expression was induced with 1 mM IPTG at OD_600_ 0.3–0.4^29^. Cells were harvested at an OD_600_ of 1.5-1.8 by centrifugation (2k x g speed, for 15 min at RT), washed two times and resuspended in 2x v/w S30A buffer (50 mM Tris base, 14 mM magnesium glutamate, 60 mM potassium glutamate, 2 mM dithiothreitol (DTT), pH 7.7). Cells were lysed by first freezing them at −70 °C then thawing the biomass and vortexing for 5 min, followed by an incubation at room temperature for 90 minutes. Following lysis, the cell extract was clarified using a small tabletop centrifuge (4 °C, 20k x g, 90 minutes) instead of an ultracentrifuge to improve accessibility. The supernatant lysates were stored in 30 μL aliquots at −70 °C for further use.

### Vacuum desiccation of cell-free expression lysates

Vacuum desiccation of *E*.*coli* lysates was performed in PCR tubes containing 10 μL lysate aliquots. Open PCR tubes were placed inside a table top vacuum desiccator containing 500 g of activated silica beads. Desiccation was conducted at room temperature and a pressure of 11 mbar over-night. After desiccation, dry aliquots were placed in plastic bags with a silica bead sachet, flushed with nitrogen and vacuum sealed using an Anova™ impulse vacuum sealer. Sealed bags were stored at room temperature in a closed styrofoam box for one week or up to a month.

### Rehydration and testing of desiccated lysate samples

All reactions were prepared in a cold room (4 °C) to minimize variations from room-temperature exposure during handling. Desiccated lysates were rehydrated to the original 10 μL volume with nuclease-free water. To ensure homogeneity and reduce mixing variations, water was added to the dried lysate, and the mixture was mixed 60 times with periodic rinsing of the tube walls to remove adherent particles. DNA, energy buffer, and MGO were premixed into an energy mix, and then combined with the rehydrated lysate samples to final master mix as described below. Reactions were then measured in the plate reader as further described. To account for experimental variability, a fresh lysate sample, of each type and same batch, was included in every run as a relative benchmark.

### Cell free reactions and dual reporter measurements

Plate reader experiments were conducted in technical triplicates of 5 µl CFES reactions. The 17 µl CFES master mix was prepared by mixing 0.64 µl plasmid DNA (3.5 nM final concentration), 2.25 µl MGO dye (30 µM final concentration; (CasNo. 2437-29-8)), 6.8 µl of *E. coli* BL21(DE3) lysate variants and 7.31 µl of energy buffer (final buffer composition in reaction mix: 7.23 mM Mg-glutamate, 72.24 mM K-glutamate, 1.55 mM amino acids solution, 51.50 mM HEPES, 1.55 mM NTPs, 0.27 mM CoA, 0.34 mM NAD, 0.77 mM cAMP, 0.07 mM folinic acid, 1.03 mM spermidine, 1.03 mM putrescine, 30.95 mM 3-PGA, 1.03 mM DTT, 10.32 mM ammonium glutamate, 4.13 mM oxalic acid, 2.06 % PEG-8000.)^37,38^. Plates were measured from the top in a Tecan Spark® microplate reader at 29 °C for 4 h. We set MG excitation to 590 nm, emission to 656 nm, with a bandwidth of 20 nm, and a signal amplification gain 80. sfGFP fluorescence was measured using an excitation wavelength of 485 nm and an emission wavelength of 535 nm with bandwidth of 20 nm and gain 30 signal amplification.

### Data analysis of fluorescence measurements

Fluorescence values were background-subtracted using negative controls (CFES reaction without template) and normalized to the fresh lysate controls. For each experiment, we calculated the normalized transcriptional and translational activity as follows: the maximum is defined as the point at which the change in fluorescence switches from positive to negative. If no such point is identified within the time-frame, the global maximum within the time frame is defined as the maximum. Normalization was performed by dividing the maximum activity of the dehydrated lysate with the equivalent of a fresh sample from the same batch. Changes in transcription (ΔMG) were quantified by calculating the difference in malachite green fluorescence between time points, as specified in figure captions. All statistical calculations and graphical visualisations were conducted using free open source Python libraries: *Numpy, Scipy, Matplotlib, Pandas* and *Seaborn*.

### *In Silico* analysis of PrCAHS1

To model the dimerisation of PrCAHS1, AlphaFold2 was used to predict monomeric and dimeric structures through the ColabFold pipeline^55^. Predictions were conducted with three recycles for monomers and 20 recycles for dimeric structures (AlphaFold2 multimer v3). The best performing model was subsequently energy minimized (relaxed) in five iterations. Predicted structures were visualized using PyMOL, where the hydrophobicity was highlighted using a Python-based script^56,57^. Sequence properties were predicted using different web-based bioinformatics tools: using the IUPRED 3 webserver, the intrinsic disorder of each residue of PrCAHS1 was predicted ^58^; charges of individual residues were calculated utilizing the Protein-Sol webserver^59^; the secondary structure identity of each residue was predicted using the PSSpred functionality of the MPI bioinformatics toolkit platform^60,61^.

### PrCAHS protein purification and SDS-PAGE

Overnight 5 mL LB cultures (37 °C, 200 rpm) from glycerol stocks were used to seed 250 mL LB cultures. At OD_600_ = 0.4, cultures were induced with 1 mM IPTG, and grown for 4 h post-induction. Cells were harvested (2k × g, 15 min, RT), resuspended to OD_600_ = 20 in HEPES buffer (50 mM HEPES, 50 mM NaCl, pH 8) with protease inhibitors (EDTA-free) tablets (Pierce™, Thermo Fisher, #A32965) and egg-white lysozyme (1 mg/mL final concentration). Resuspensions were incubated with lysozyme at 5 mL volume in 15 mL tubes (37 °C, 200 rpm, 60 min), sonicated (40 % amplitude, 10 s on/off, 4 min total), and centrifuged at high speeds (20k × g, 90 min) to mimic conditions during lysate production. Supernatants were stored on ice, while pellets with insoluble protein (showed in Fig. 4) were washed (4-6 mL/g biomass) two times with HEPES buffer (50 mM HEPES, 100 mM NaCl, pH = 8) and resolubilised through boiling at 95 °C (90 minutes). For earlier purifications such as in Fig. S3, pellets were washed in PBS buffer (137 mM NaCl, 2.7 mM KCl, 10 mM NaHPO_4_*12 H_2_O, 2 mM KH_2_PO_4_), freeze-thawed (−70 °C; RT), and finally resuspended in PBS + 2 M urea buffer (1/3 original lysis reaction volume), based on previous works^62–65^. Independent of the solubilisation method used, final centrifugation was done at 20k × g for 10 minutes. Proteins were concentrated further using Amicon® Ultra-filter columns (Sigma-Aldrich) using the manufacturer’s protocol.

Purified protein and lysate samples were analysed by SDS-PAGE. For lysate samples, 30 ul aliquotes were boiled at 95°C for 30 minutes and centrifuged as described above prior processing for visualization. 1 to 5 μL of each sample was supplemented with 5 μL of 4x NuPage LDS PAGE sample buffer (ThermoScentific), 1 μL of β-mercapto-ethanol and filled up with PBS buffer to 20 μL. Following a heat treatment at 70 °C for 10 min the samples and a protein standard (PageRuler™, Thermofischer Scientific) were loaded onto the SDS-PAGE (12 % SDS-Tris-glycine) gels.

### Characterisation of TFE-induced PrCAHS assemblies

For turbidity measurements and microscopy of purified proteins, 20 % and 15 % of 2,2,2-trifluoroethanol (v/v) respectively, was used. TFE was mixed with either purified PrCAHS proteins or BSA protein (10 mg/mL final protein concentration used). Samples were incubated for 1 hour at 4 °C prior to visualisation and turbidity measurements using Implen NanoPhotometer at OD_600_. PrCAHS proteins were redissolved through dilution in nuclease-free water (1:4), followed by a subsequent heating step at 95 °C for 10 minutes.

Further imaging was carried out by pipetting 5 µl of reaction mixes (incubated with 15 % TFE) on Lumox® dishes (Sarstedt AG & Co KG). Plates were transferred to a Leica DMI8 inverted fluorescence microscope and imaged in the brightfield at 20 x magnification.

## Supporting Information

Figure S1: Current model of CAHS protein assembly into higher order structures under desiccation

Figure S2: Plasmid maps of constructed vectors

Figure S3: 12% SDS Page assessing heterologous CAHS protein production in *E. coli* BL21(DE3)

Figure S4: Workflow for the low-cost desiccation of cell-free expression lysates

Figure S5: Normalized maximum sfGFP production in individual short term desiccation experiments

Figure S6: MG kinetics from individual short term desiccation experiments grouped by damage

Figure S7: Rate of change in MG signal kinetics

Figure S8: Workflow for the purification of CAHS proteins

Figure S9: *In Silico* modelling of PrCAHS1

Table S1: Plasmids used in this study

## Authors information

### Notes

The authors declare no competing financial interest.

### Author contributions

MM: Conceptualization, Methodology, Investigation, Visualization, Writing – original draft, Writing – review & editing. IB: Conceptualization, Methodology, Visualization, Supervision, Writing – review & editing. AG: Conceptualization, Methodology, Visualization, Supervision, Writing – review & editing. HN: Conceptualization, Funding acquisition, Supervision, Writing – review & editing.

## Acknowledgments

This work was funded by the Deutsche Forschungsgemeinschaft (DFG, German Research Foundation) grant NI 2040/1-1 and the European Research Council (ERC Starting Grant SYNSEMBL, 101078028). MM acknowledges support by the Studienstiftung des Deutschen Volkes scholarship program, which provided financial support during his studies and work on this publication. We thank the iGEM Straubing 2024 team members who worked on the initial stages of this idea; we also thank Emma Crean and Jan Kalkowski (TUM) for helping to supervise the iGEM project; Anibal Arce (Northwestern University) and Prof. Dr. Fernando Guzman-Chavez (UNAM) for scientific input during the iGEM project; Tommi Bui and Prof. Dr. Wenwen Fang (TUM) for helpful discussions about protein gels and material properties; and finally Han Banlaki-Kok for inspiring this project. Schematics were generated using Biorender.com.

## Abbreviations

CFES: Cell-free expression system
CAHS: tardigrade-derived cytosolic-abundant heat-soluble protein
EL: standard lysate produced from a strain carrying the autolysis plasmid plus the empty expression vector
L: standard lysate produced from a strain containing only the autolysis plasmid
MG: Malachite green
MGO: Malachite green oxalate
PrCAHS: *Paramacrobiotus richtersi*-derived cytosolic-abundant heat-soluble tardigrade proteins
sfGFP: superfolder green fluorescence protein
TFE: 2,2,2-trifluoroethanol
TL: tardigrade lysate produced from a strain expressing PrCAHS1 protein

