## Supplementary Information for "Tardigrade-Derived Strategy for Low-Cost Storage of Cell-Free Expression Lysates"

*Shared first authorship

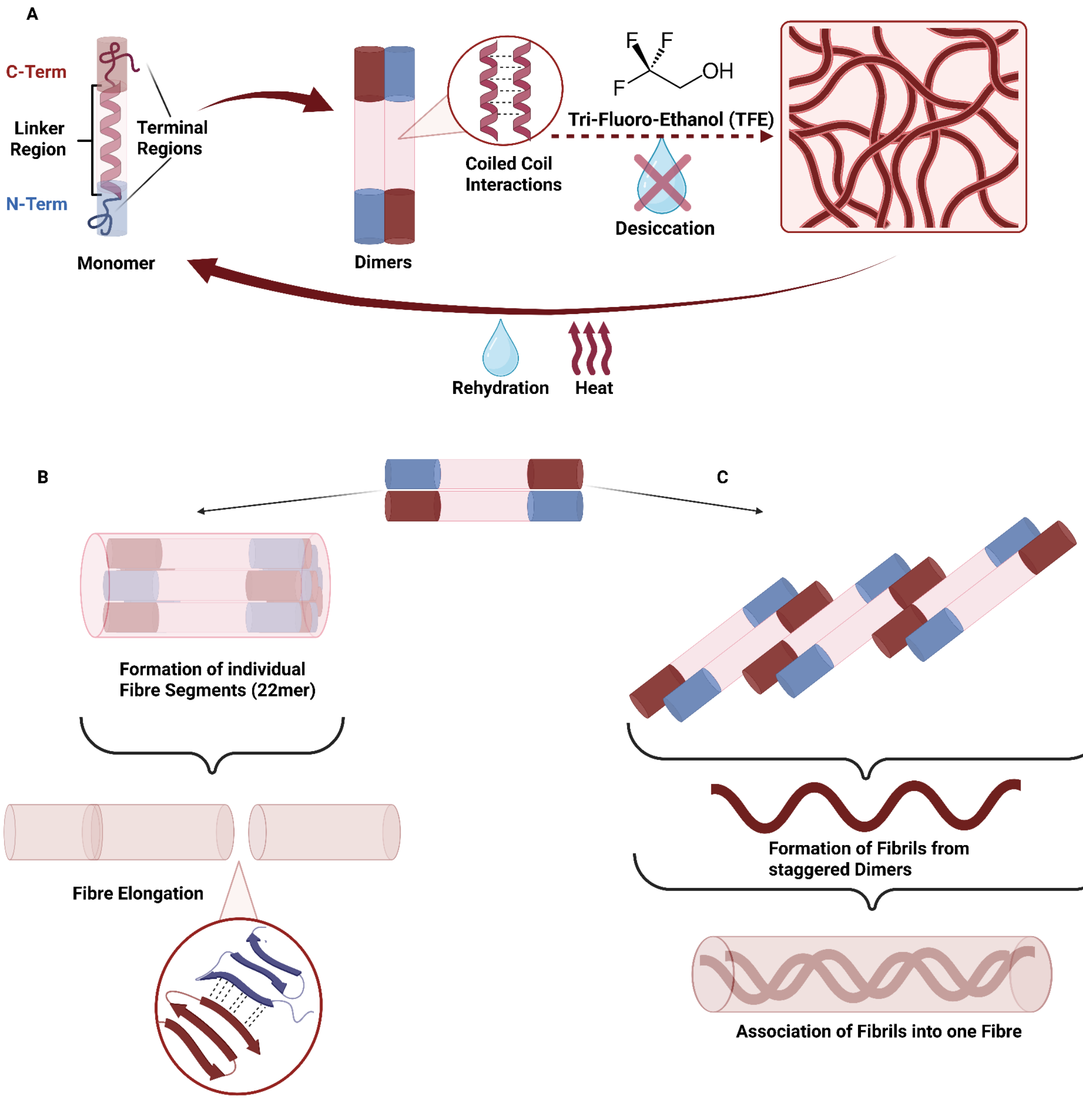


**Figure S1. Current models of CAHS protein assembly into higher order structures during desiccation**
**(A)** Simplified visualisation of a CAHS monomer, highlighting essential structural features. Antiparallel association of monomers into dimers, driven by coiled coil interaction of the linker regions, and further assembly into fibrous networks during desiccation[1]. **(B)** Proposed oligomerisation of dimers into 22-mer fibre bundles and elongation of fibres mediated through ß-ß interactions of "sticky" terminal regions[1]. **(C)** Alternative model, proposing polymerisation of dimers into fibrils which associate to form higher order fibres[2].

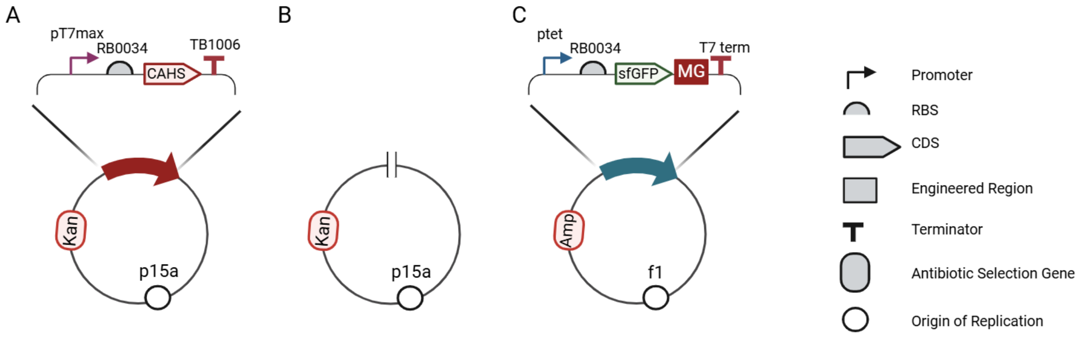


**Figure S2. Plasmid maps of constructed vectors**
**(A)** Vector used for the expression of CAHS proteins (used for lysate generation as well as protein purification) **(B)** Empty Vector used for the generation of a control Lysate lacking the CAHS protein. **(C)** Plasmid used as the DNA template coding for the sfGFP translational reporter and malachite green aptamer transcriptional reporter used in the assessment of lysate activity. RBS = Ribosome Binding Site, CDS = Coding Sequence , MG = Malachite Green , Kan = Kanamycin , Amp = Ampicilin.

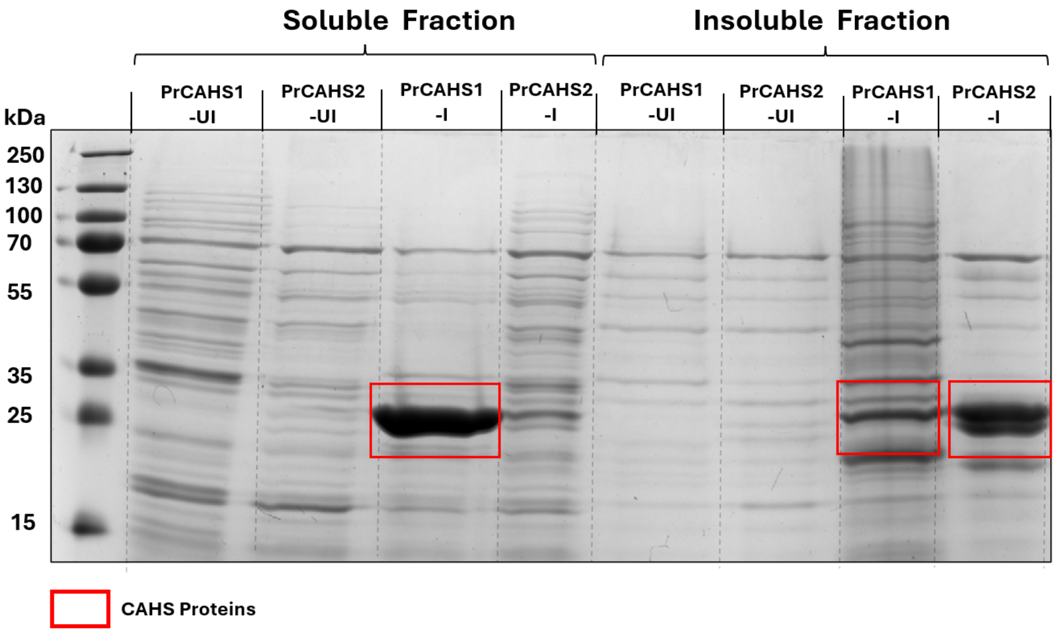


**Figure S3. 12% SDS Page assessing heterologous CAHS protein production in *E. coli* BL21(DE3)**
Lanes: (1) PageRuler™ prestained protein ladder (Thermofischer Scientific); (2-5) Protein Extracts from soluble fraction (5 µl supernatant following lysis); (6-9) Protein samples (5 µl) generated from insoluble fractions (Pellet following lysis, resolubilized using urea (see Methods)). (2-3,6-7) Protein samples (5 µl) taken immediately prior to induction at OD600=0.2 (4-5,8-9) Samples taken following four hours of induction (1mM IPTG) (5 µl). Abbreviations: PrCAHS1 = Samples taken from *E. coli* BL21(DE3) cultures harboring the PrCAHS1 expression vector, PrCAHS2 = Samples taken from *E. coli* BL21(DE3) cultures harboring the PrCAHS2 expression vector , UI = uninduced, I = Induced Expected size: PrCAHS1 = 26.5 kDa, PrCAHS2 = 25.5 kDa

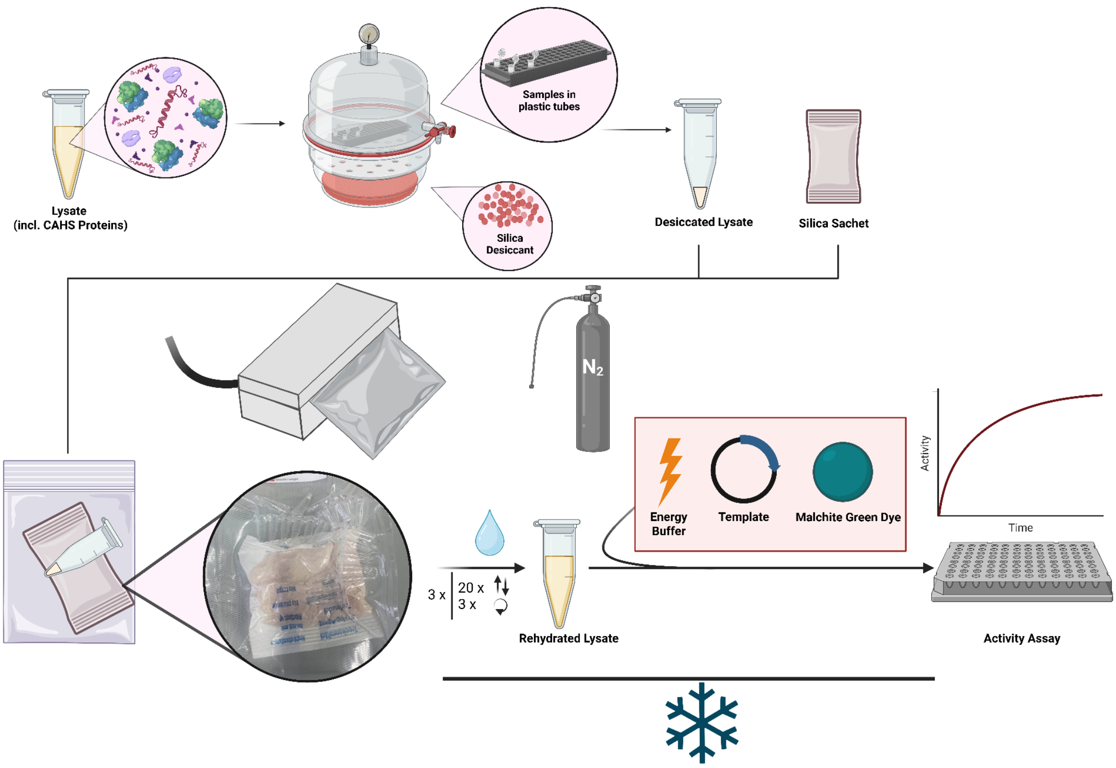


**Figure S4. Workflow for the low-cost desiccation of cell-free expression lysates**
Open tubes of lysates containing CAHS protein (or controls) were placed inside a conventional vacuum dessicator containing silica dessicant. Following dessication at 11 mbar, at room temperature, the dessicated samples were placed inside a plastic bag with a silica sachet, flushed with liquid nitrogen, and subsequently vacuum sealed utilizing an impulse vacuum sealer. After storage at room temperature, samples were rehydrated, in a cold room, to the original volume using deionized water and mixed thoroughly as indicated. The rehydrated lysate is supplemented with corresponding volumes of energy buffer, plasmid DNA template and malachite green dye and finally transferred into a plate reader where the activity is measured.

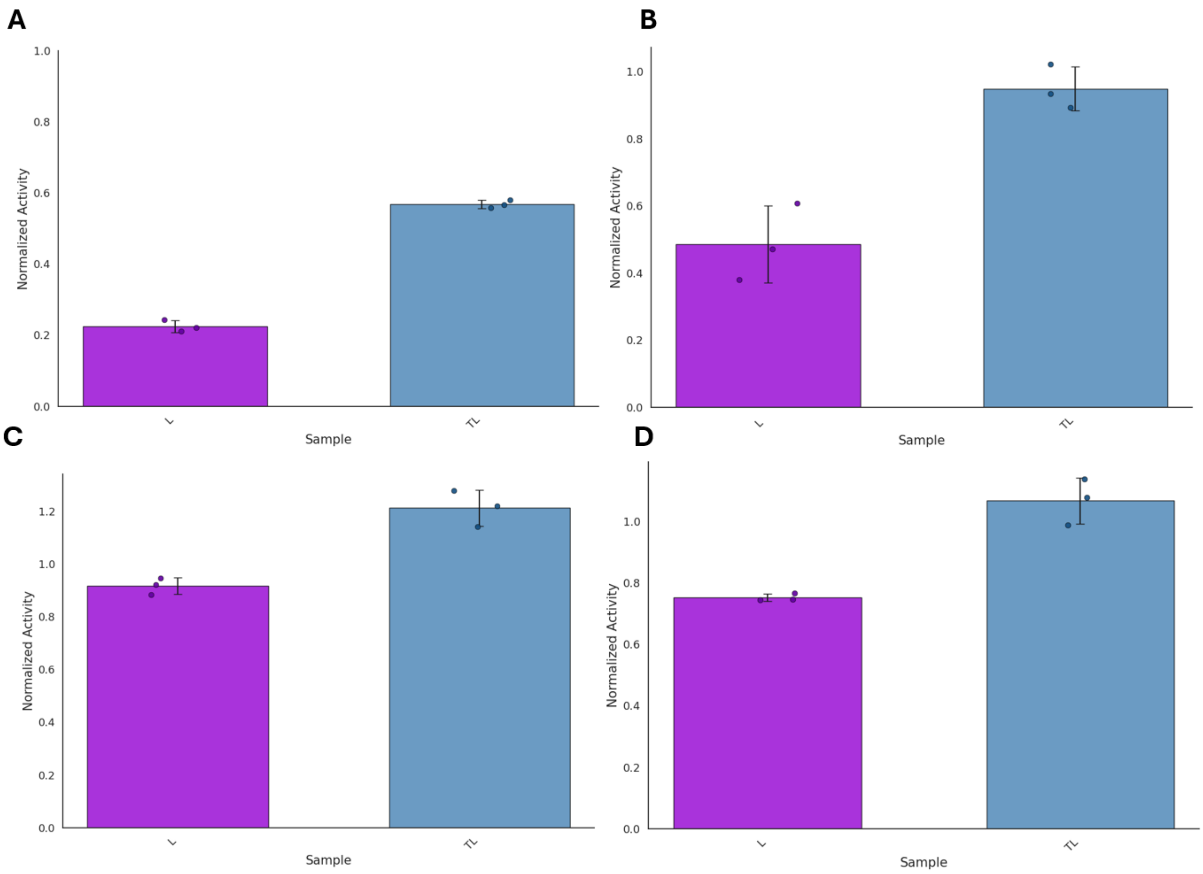


**Figure S5. Normalized maximum sfGFP production in individual short term desiccation experiments**
**(A-D)** Maximum normalized sfGFP concentrations of different lysate types L (purple) and TL (blue) after 1 week of desiccation. Bars show means ± SD (n = 3). Individual data points are overlaid. **(A-D)** are the results from separate desiccation experiments in Fig. 2B (01-04).

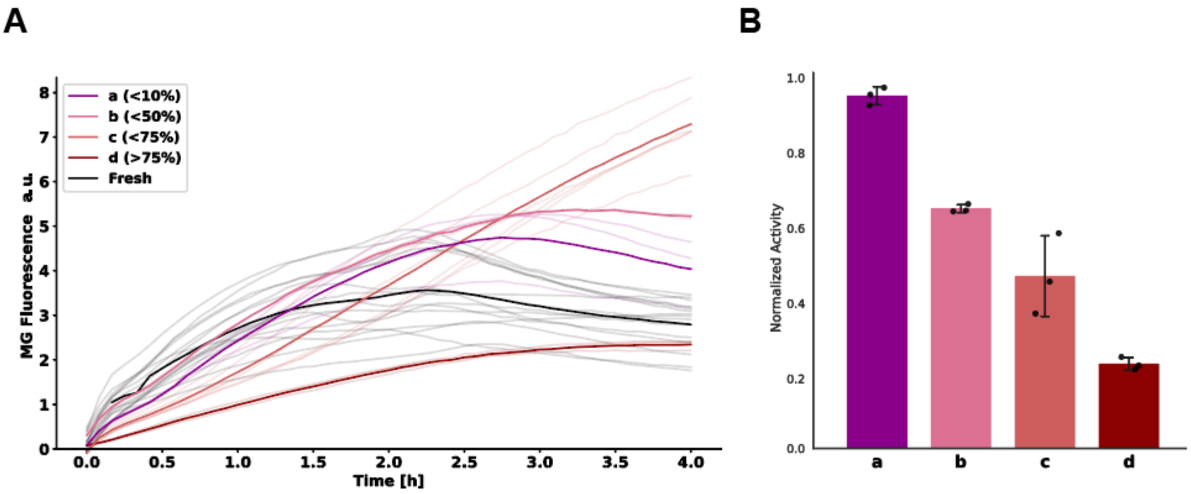


**Figure S6. MG kinetics from individual short term desiccation experiments grouped by damage**
**(A)** Malachite Green (MG) fluorescence kinetics across four separate short term (1 week) desiccation experiments of standard lysate not containing tardigrade CAHS proteins. Solid lines represent means , faint traces represent individual replicates. Colors and letters denote damage categories defined by the inverse of normalized sfGFP concentration from the corresponding samples. (a) <10% damage, (b) 10–50%, (c) 50–75%, and (d) >75%. **(B)** Normalized activity of rehydrated standard lysates calculated from endpoint sfGFP signal.

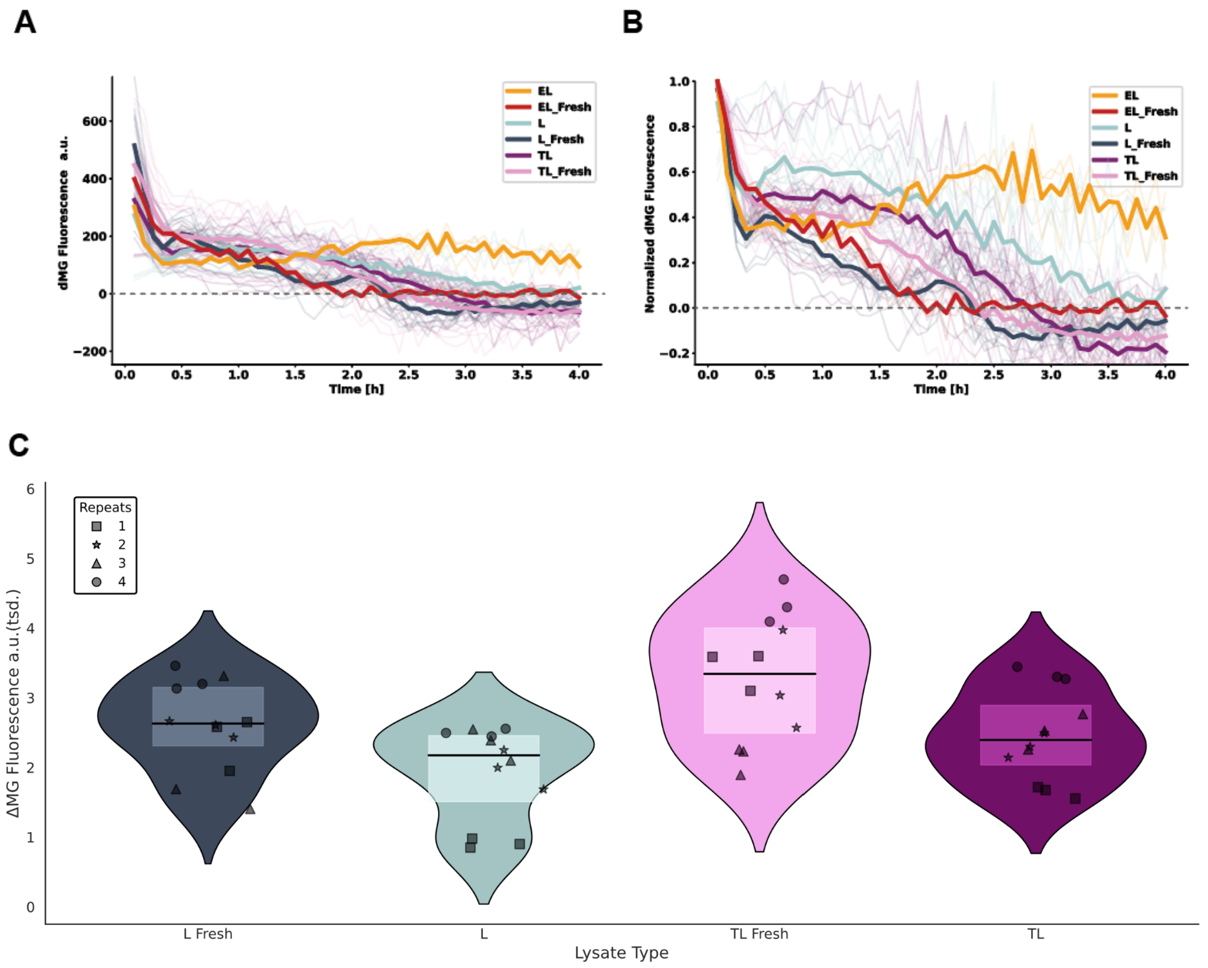


**Figure S7 Rate of change in MG signal kinetics**

**(A)** ΔMG kinetics of different lysate types and treatment. Bold lines are mean values of replicates. The dashed, grey line marks the inversion where the MG signal started to decrease over time. **(B)** Normalized ΔMG kinetics emphasize the different trends over time between fresh, intact and degraded lysates. Running ΔMG kinetics were calculated by subtracting MG fluorescence at time point n (n times 5 minutes) from the preceding value at time point n-1. **(C)** Initial mRNA synthesis rates. Violin plot comparing ΔMG between 1h and 0h across four separate 1-week desiccation experiments. Symbols distinguish individual experiments (n = 12). The line within the violin plot represents the median, upper and lower border of the box plots representing upper and lower quartiles respectively.

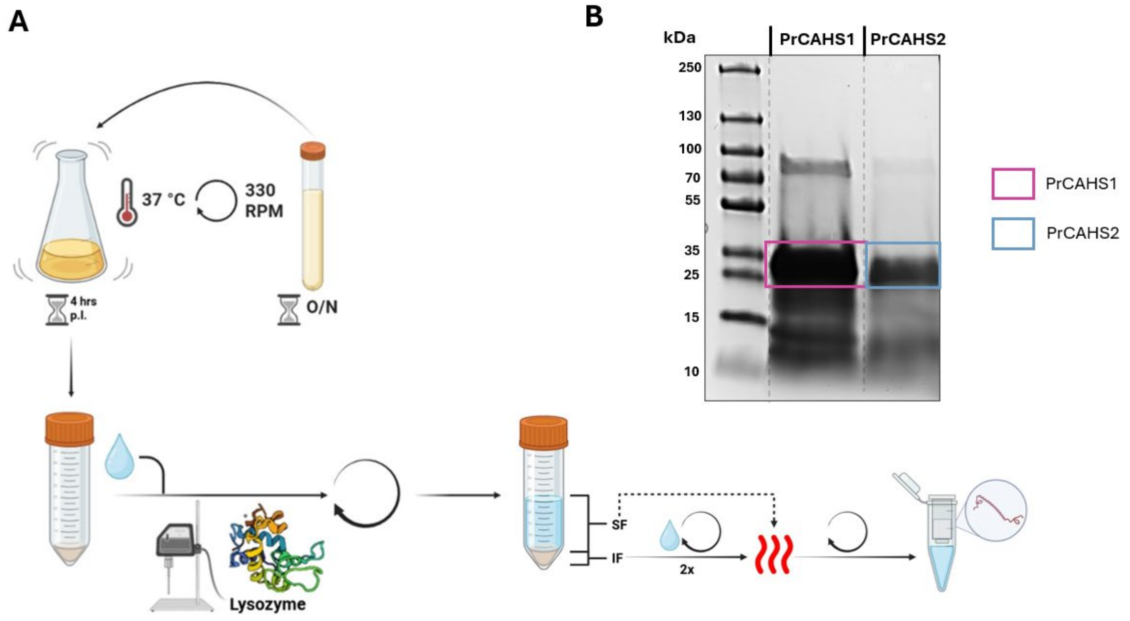


**Figure S8. Workflow for the purification of CAHS proteins**

**(A)** Schematic of the CAHS protein purification workflow. Overnight cultures (5 mL) of *E. coli* BL21(DE3) cells carrying the CAHS expression construct (table S1) were used to inoculate 250 mL LB cultures. Following induction at $OD_{600}$ = 0.3–0.4, cells were incubated for 4 h before biomass collection. Pelleted cells were resuspended in buffer (composition variable) and lysed by combined lysozyme treatment and ultrasonication. Soluble and insoluble fractions were separated by centrifugation. After multiple washes of the insoluble fraction, both soluble and insoluble fractions were subjected to heat treatment (95 °C, ≤90 min). CAHS proteins remained in the soluble fraction after heat treatment and were further concentrated using Amicon® filter units. **(B)** Exemplary SDS page (12 %) showing the CAHS proteins extracted from *E. coli* BL21(DE3) using the established purification protocol. (5 µl of samples were loaded); Lanes: (1) PageRuler™ prestained protein ladder (Thermofischer Scientific) (2) PrCAHS1 (5 µl), (3) PrCAHS2 (5 µl). Expected size: PrCAHS1 = 26.5 kDa, PrCAHS2 = 25.5 kDa

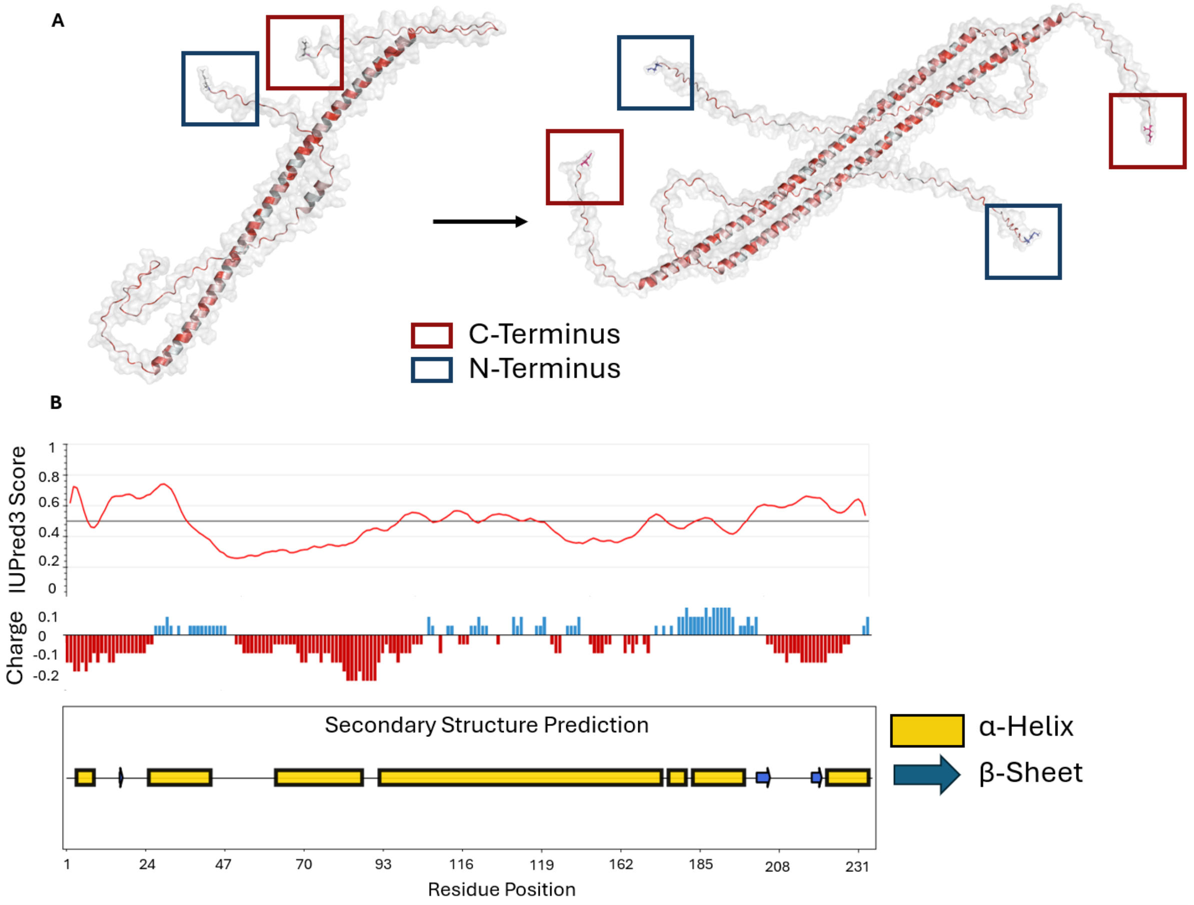


**Figure S9. In Silico modelling of PrCAHS1**

***(A)*** PyMOL visualisation of AlphaFold2 predictions of PrCAHS1 secondary structure (monomeric) and in a dimeric state. Termini are indicated with blue (N-Terminus) and red (C-terminus) respectively. Individual residues were colored in a hydrophobic gradient: non-hydrophobic (white) to hydrophobic (red) amino acids[3].

**(B)** Sequence property predictions. Top: intrinsic disorder of residues predicted using IUPRED 3 webserver[4]. Middle: Charge of individual residues estimated using the Protein-Sol software[5]. Bottom: Secondary Structure prediction of PrCAHS1 using the PSSpred functionality of the MPI bioinformatics toolkit platform[6,7].

**Table S1.** Plasmids used in this study; promoter RBS gene terminator, gene sequences are capitalised

| **Plasmid** | **Description** | **Reference** | **Relevant sequence** |
|---|---|---|---|
| pCAHS1 | Plasmid used for the heterologous expression of PrCAHS1 for purification and lysate generation.<br><br>Components: pJUMP26-1A(sfGFP)_p**T7max**::**RB0034**::**PrCAHS1**::**TB1006**, Kanamycin Resistance, p15a | this study | **gcctttcgttttatttgatgcctttaattaaggagttttgcaggtgccttggaacacctgcttttcgctgaattcgcggccgcttctagagcgtctctggagAATTCTAATACGACTCACTATAGGGATACTAGAGAAAGAGGAGAAATAATCAATGTCAGCGGAAGCTATGAACATGAACATGAACCAGGATGCAGTTTTTATTCCTCCTCCGGAAGGCGAACAGTATGAACGCAAAGAGAAACAAGAGATTCAACAGACGTCGTACCTGCAAAGCCAGGTAAAGGTTCCGCTGGTCAACCTGCCCCGCACCTTTCTTCTCAACCTCCTTCTCCGCGCAGGAGATCCTTGGGGAAGGTTTCCAGGCTAGTATTTCTCGGATTTCCGCCGTTAGTGAAGAACTGTCCTCTATCGAAATTCCGGAGCTGGCGGAAGAGGCCCGTCGTGACTTTGCGGCCAAAACGCGCGAACAGGAGATGCTGTCTGCTAACTATCAAAAAGAAGTTGAGCGCAAAACTGAAGCCTATCGGAAGCAGCAAGAAGTGGAGGCAGACAAAATTCGGAAAGAGCTGGAAAAGCAGCATCTGCGTGACGTGGAGTTCCGCAAAGATATTGTAGAGATGGCAATCGAAAACCAGAAAAAAAATGATTGATGTCGAAAGCCGTTATGCAAAAAAAAGATATGGACCGCGAACGTGTGAAAGTCCGTATGATGCTCGAGCAGCAAAAATTTCATAGCGACATCCAGGTAAATCTCGATTCTAGCGCAGCAGGGACCGAAACTGGAGGCCAAGTTGTTTCAGAATCTCAGAAATTTACCGAACGTAATCGTCAGATTAAACAGGCTTAAAAAAAAAAAACCCCGCCCCTGACAGGGCGGGGTTTTTTTTcgctaatgtgagacgtactagtagcggccgctgcagggagttgtcttcgaagacttcgctctagtcttggactc** |
| pCAHS2 | Plasmid used for the heterologous expression of PrCAHS2 for purification.<br><br>Components: pJUMP26-1A(sfGFP)_p**T7max**::**RB0034**::**PrCAHS2**::**TB1006**, Kanamycin Resistance, p15a | this study | **gcctttcgttttatttgatgcctttaattaaggagttttgcaggtgccttggaacacctgcttttcgctgaattcgcggccgcttctagagcgtctctggagAATTCTAATACGACTCACTATAGGGATACTAGAGAAAGAGGAGAAATAATCAATGGAGGCCATGAATATGAATATCCCCCGCGATGCCATGTTTGTTCCGCCACCGGAATCTGAGCAAAATGGGTATCATGAGAAGTCAGAAGTTCAGCAAACAAGTTATATGCAGAGTCAAGTCAAAGTGCCACATTATAATTTCCCGACACCATATTTTACGACTTCCTTTTCTGCGCAAGAGCTGCTGGGCGAAGGGTTTCAAGCCTCAATTTCCCGTATTTCAGCCGTTACGGAAGACATGCAGAGCATGGAAATCCCGGAGTTCGTTGAAGAGGCCCGCCGTGATTACGCAGCCAAAACACGTGAAAATGAGATGCTGGGGCAACAATATGAAAAAGAGCTGGAACGTAAGTCCGAAGCCTACCGCAAACATCAGGAAGTAGAGGCCGACAAAATCCGCAAAGAACTTGAAAAACAGCATATGCGTGATATTGAATTTCGGAAAGAAATTGCAGAACTGGCGATTGAGAACCAAAAACGTATGATCGATCTTGAATGCCGCTATGCAAAAAAAGACATGGACCGGGAACGCACAAAAGTTCGTATGATGCTGGAGCAACAGAAATTCCATAGTGATATCCAGGTAAATCTGGATTCTTCTGCGGCTGGGACCGAGAGCGGAGGTCATGTAGTGAGCCAGTCTGAAAAGTTCACCGAACGTAACCGCGAGATGAAACGCGCTTAAAAAAAAAAAACCCCGCCCCTGACAGGGCGGGGTTTTTTTTcgctaatgtgagacgtactagtagcggccgctgcagggagttgtcttcgaagacttcgctctagtcttggactcctgttgatag** |
| pEmpty | Plasmid used for the generation of the empty vector control lysate.<br><br>Components: pJUMP26-1A(sfGFP), Kanamycin Resistance, p15a | this study | |

| | | | |
|---|---|---|---|
| psfGFP-MG | Dual reporter plasmid to evaluate effects of desiccation on transcription and translation<br><br>Components:<br>MC_2-5-DO**_pTet::RBS**::**sfGFP**::3xSTOP::**MG**::**T7terminator**<br>Ampicilin Resistance ,ColE1 | this study | **ggagaggcggttgtagtgggggaatggatagcaagctgcgggtagac<br>ctccaattattgaaggcctcccaaatcggggggcctttttattgataacaaa<br>aGGAGCTGTTCTCAGTGATAGAGATTGACATCCCTAT<br>CAGTGATAGATATAATGAGCACTACTAGAGAGAAGGA<br>GGAAAAAAAAATGCGTAAAGGTGAAGAACTGTTTACC<br>GGTGTTGTTCCAATTCTGGTTGAACTGGATGGTGATGT<br>TAACGGTCACAAATTTTCTGTTCGTGGTGAAGGCGAA<br>GGTGATGCAACCAACGGTAAACTGACCCTGAAATTTA<br>TCTGTACCACTGGTAAACTGCCAGTTCCATGGCCAAC<br>TCTGGTTACCACTCTGACCTACGGTGTTCAATGTTTTG<br>CACGTTACCCAGATCACATGAAACAACACGATTTTTT<br>CAAAAGCGCAATGCCAGAAGGTTACGTTCAAGAACG<br>TACCATCTCTTTTAAAGATGACGGCACCTACAAAACC<br>CGTGCGGAAGTTAAATTTGAAGGTGATACCCTGGTTA<br>ACCGCATTGAACTGAAAGGCATCGATTTTAAAGAAGA<br>TGGTAACATCCTGGGCCACAAACTGGAATACAACTTT<br>AACTCTCACAACGTGTACATCACCGCAGACAAACAAA<br>AAAACGGTATCAAAGCGAACTTCAAGATCCGTCACAA<br>CGTTGAAGATGGTTCTGTTCAACTGGCAGATCACTAC<br>CAACAAAACACCCCCAATTGGTGATGGTCCAGTTCTGC<br>TGCCAGATAACCACTACCTGTCTACCCAAAGCGTTCT<br>GTCTAAAGATCCAAACGAAAAACGTGATCACATGGTG<br>CTGCTGGAATTTGTTACCGCAGCAGGTATTACCCACG<br>GTATGGATGAACTGTACAAAGCAGCTTTATGATGATG<br>AGGGTATGCCTGGCGACCATAGCGATTGGGTAACCG<br>GATCCCGACTGGCGAGAGCCAGGTAACGAATGGATC<br>CGGTAACCAATTAGCGCCGATGGTAGTGTGGGGTTTC<br>CCCATGTGAGAGTAGGACATCGCCAGGCATTAGCATA<br>ACCCCTTGGGGCCTCTAAACGGGTCTTGAGGGGTTTT<br>TTGcgctggacccgcgtgtcttcggagaaccatctcgtaagaggatagt<br>agttactggagacga** |
| pADLysR | Used for facile autolysis during lysate generation<br><br>Components:<br>pPro24_beta-lactamase phage lambda gene R | [8] | See Addgene [# 99244] |
